# Drug Resistance without a cost? Common and uncommon routes to fosfomycin resistance in Uropathogenic *Escherichia coli*

**DOI:** 10.1101/2023.06.20.545750

**Authors:** Tomas A. Bermudez, John R. Brannon, Neha Dudipala, Seth Reasoner, Grace Morales, Michelle Wiebe, Mia Cecala, Connor Beebout, Omar Amir, Maria Hadjifrangiskou

## Abstract

Fosfomycin kills bacteria by blocking the binding of phosphoenolpyruvate (PEP) to the bacterial enzyme MurA and halting peptidoglycan synthesis. While its use has increased with the emergence of antibiotic resistance, the mechanisms leading to fosfomycin resistance remain relatively unexplored. In uropathogenic *Escherichia coli* (UPEC) that accounts for >75% of urinary tract infections (UTIs), fosfomycin enters the cell primarily through UhpT, which transports glucose-6-phosphate (G6P) glycolysis intermediate into the cell. Mutations in *uhp*T lead to fosfomycin resistance and have been identified during antimicrobial susceptibility testing (AST) in non-susceptible inner colonies that form within the zone of inhibition. However, EUCAST and CLSI guidelines differ in how to read fosfomycin AST when such resistant colonies arise. Work from our lab and others demonstrated that glycolysis is dispensable during acute UTI. Moreover, G6P is scarce in urine, prompting us to test the hypothesis that *uhp* mutations may not impart a fitness cost to the pathogen. We report that loss of *uhp* indeed does not impair UPEC pathogenesis and that clinical isolates exist that lack the *uhp* locus altogether. Analysis of non-susceptible inner colonies revealed a suite of novel genes involved in fosfomycin resistance. One of them is PykF that converts PEP to pyruvate during glycolysis. Single deletions of *pykF* or its anaerobic homolog *pykA* do not attenuate UPEC. Based on our data, we raise the alarm that multiple routes lead to fosfomycin resistance and do not affect pathogenesis and propose that the current EUCAST and CLSI guidelines unify into how they evaluate fosfomycin AST.

**IMPORTANCE:** While fosfomycin resistance is rare, the observation of non-susceptible subpopulations among clinical *Escherichia coli* isolates is a common phenomenon during antimicrobial susceptibility testing (AST) in American and European clinical labs. Previous evidence suggests that mutations eliciting this phenotype are of high biological cost to the pathogen during infection, leading to current recommendations of neglecting non-susceptible colonies during AST. Here we report that the most common route to fosfomycin resistance, as well as novel routes described in this work do not impair virulence in uropathogenic *E. coli*, the major cause of urinary tract infections, suggesting a re-evaluation of current susceptibility guidelines is warranted.

## INTRODUCTION

Urinary tract infections (UTIs), which are among the most common bacterial infections worldwide, are a major driver of antibiotic prescription in primary care [1-2]. Uropathogenic *Escherichia coli* (UPEC) accounts for >75% UTIs, with multi-drug resistant sequence types (ST) like ST131 spreading globally [3-4]. Current first-line treatments for uncomplicated UTI include trimethoprim-sulfamethoxazole or ciprofloxacin. However, clinics experience resistance rates as high as 17.4% and 12.1% respectively for these commonly prescribed antibiotics in the United States [5]. Additionally, resistance rates in developing countries are typically much higher due to less regulation over antibiotic use [6]. Increasing trends in antibiotic resistance have led to more frequent use of the antibiotic fosfomycin, in both European and American health care facilities [7-8]. While fosfomycin prescriptions remain low compared to prescriptions for TMP-SMZ, ciprofloxacin or nitrofurantoin [9], one can predict that fosfomycin prescriptions will continue to rise in the next decade.

Fosfomycin has broad spectrum activity against both Gram-positive and Gram-negative organisms and accumulates at clinically relevant concentrations in the urinary tract for several days following a single oral administration [10]. Fosfomycin enters the bacterial cell through at least two sugar transporters, UhpT (**Fig. 1A**) and GlpT, which transport glucose-6-phosphate (G6P) and glycerol-3-phosphate (G3P) respectively [11]. This is believed to be due to structural similarities between fosfomycin and their cognate targets (**Fig 1C**). Interfering with the first committal step of peptidoglycan synthesis, fosfomycin acts as a phosphoenolpyruvate (PEP) analog, irreversibly binding to the enoylpyruvate transferase MurA (**Fig. 1B**) and leading to cell lysis [12]. Expression of the *uhpT* transporter is regulated by the UhpBAC signaling system (**Fig. 1A**) [13] and loss-of-function mutations in *uhpBAC* or *uhpT* genes have been associated with increased resistance to fosfomycin [14]. Non-susceptible inner colonies (NICs) that commonly arise on agar plates during Kirby-Bauer antimicrobial susceptibility testing (AST) of clinical isolates, most frequently harbor mutations in *uhpT* or *glpT* [14], [15].

**Fig. 1.**
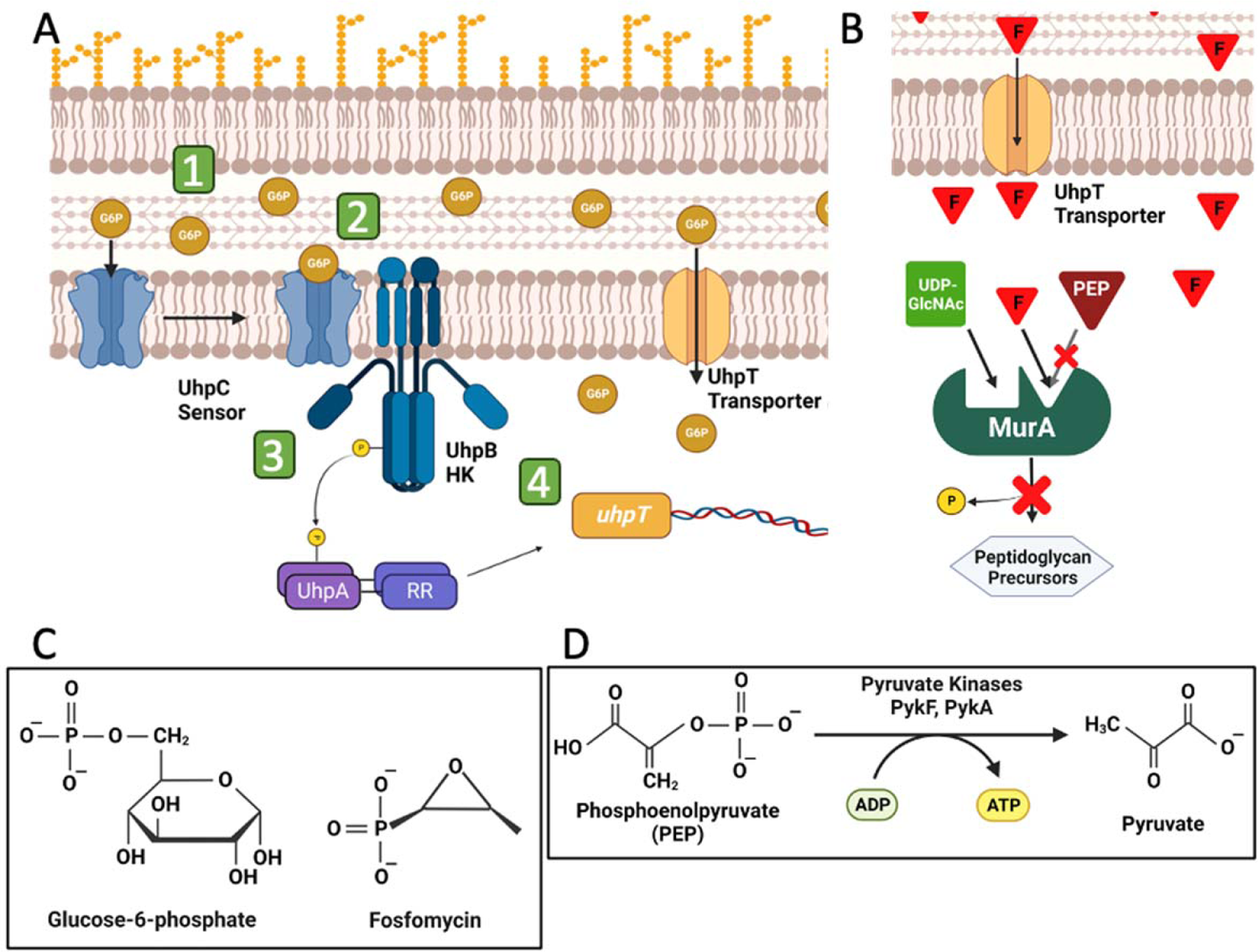
The UhpBAC two-component system detects and regulates the importation of Glucose-6-Phosphate (G6P). **A)** Schematic depicts the UhpC-BA signaling cascade as it has been delineated for K12 *E. coli*: (1) UhpC transmembrane sensor binds G6P. (2) UhpC-G6P interacts with UhpB histidine kinase (HK), stimulating UhpB autophosphorylation. (3) UhpB-P serves as a phosphodonor for the UhpA response regulator (RR). (4) UhpA-P upregulates *uhpT* expression, increasing G6P import. **B)** Fosfomycin acts as a phosphoenolpyruvate (PEP) analog, irreversibly binding the enoylpyruvate transferase MurA in place of PEP, restricting peptidoglycan synthesis. **C)** Fosfomycin shares some structural similarities with G6P, which presumably facilitate its entry into the cytoplasm through the UhpT transporter. **D)** The pyruvate kinases PykA and PykF catalyze the transfer of the phosphate group of PEP to an ADP molecule, forming pyruvate and ATP.

Previous studies concluded that fosfomycin resistant subpopulations rarely arise *in vivo*, because the inability to import G3P or G6P would be accompanied by a high biological fitness cost to the pathogen [16-17]. However, UPEC does not rely on glycolysis during acute UTI [18-19] nor is there free G6P that is abundant in the urine [20]. These observations prompted us to carefully evaluate how emergence of fosfomycin resistance may influence UPEC fitness during infection and during asymptomatic colonization of reservoir niches. In this work, we leverage a large, representative cohort of UPEC isolates from Vanderbilt University Medical Center’s *micro*VU repository to evaluate the frequency of fosfomycin resistance (Fos^R^) arising *in vitro*; define new mutations associated with Fos^R^ and determine the impact of identified Fos^R^-associated mutations to UPEC fitness. We report that >70% of UPEC readily form Fos^R^ subpopulations *in vitro* and that indeed most mutations associated with fosfomycin resistance are within the UhpBAC locus, or its regulated target *uhpT*. We go on to demonstrate using a well-established murine model of UTI that loss of *uhpBAC*, or *uhpT* do not impart a fitness cost to UPEC, during acute, or chronic infection. Moreover, we demonstrate the corresponding *uhp* mutants demonstrate no defects in the formation of gut reservoirs, indicating that UPEC does not rely on G6P uptake during acute infection of the bladder or during chronic colonization of the gut. We show that ∼ 35% of fosfomycin resistant subpopulations tested in our studies do not harbor mutations in *uhp* genes, pointing towards additional yet uncharacterized routes to fosfomycin resistance. Of these newly identified mutations imparting fosfomycin resistance, we focused on *pykF*, the aerobic pyruvate kinase that converts PEP to pyruvate and ATP (**Fig. 1D**). Clean deletion of *pykF* in UPEC to increased fosfomycin resistance, presumably because the loss of this enzyme increases PEP levels for outcompeting fosfomycin binding to MurA. Deletion of the anaerobic pyruvate kinase gene, *pykA,* does not affect fosfomycin resistance, but the double deletion imparts an additive increase. In the urinary tract (infection niche) and the gut, neither of the single deletion mutants loses fitness over time, while the double mutant does display a fitness defect during infection, indicating that PykA and PykF exhibit functional redundancy in the host.

Collectively, this work demonstrates that not all mutations associated with resistance are accompanied by a fitness cost, raising awareness in initiating the efforts to curb future increases in fosfomycin resistance. Moreover, our work suggests that the formation of NICs during fosfomycin AST should not be ignored.

## RESULTS

### Analysis of fosfomycin resistant (Fos^R^) subpopulations arising during clinical antimicrobial susceptibility testing (AST)

Recent clinical screenings indicate that urinary *E. coli* isolates have maintained rates of fosfomycin resistance as low as 3.6% [21]. However, fosfomycin AST is not performed routinely in the clinical lab and, when performed, a common observation during Kirby-Bauer susceptibility testing is the occurrence of resistant subpopulations that grow within the parental zone of inhibition (**Fig 2A, C** and [22]). We thus wanted to evaluate the prevalence of this phenomenon in UPEC isolates. Vanderbilt’s *micro*VU repository banks over 13,000 UPEC isolates yearly. To accurately sample this cohort a power analysis was conducted, determining that screening of 329 isolates would be sufficient to estimate the average fosfomycin non-susceptible inner colony (NIC) count of the 13,000 *micro*VU annual isolates with a margin of error of 5 and a confidence interval of 95%. Subsequent screening of 337 clinical strains (**Supplementary File 1**) from UTI patients seen at Vanderbilt University Medical Center demonstrated that in 77% of the tested strains fosfomycin NICs are observed during susceptibility testing, with NIC numbers ranging from 5 to 50+ per plate (**Fig. 2B, Supplementary File 1**). Overall, NIC count did not correlate to zone of inhibition size following EUCAST guidelines, and tested isolates demonstrated on average 27.8 NICs per disk diffusion plate (**Fig. 2B, Supplementary File 1**). The UPEC isolates used in our study were acquired from the Vanderbilt *micro*VU biorepository, where each clinical isolate is linked to the de-identified clinical record of the source patient. Using this resource, we asked if the number of NICs produced by UPEC strains correlate with specific disease outcome or pre-existing patient condition, like type 2 diabetes mellitus [23]. We found that the number of NICs per isolate do not correlate to diabetic status (**Fig. S1A, Supplementary File 2**), UTI symptomology (**Fig. S1B, Supplementary File 2**) or patient age (**Fig. S1C, Supplementary File 2**). These data suggest that there is no correlation between clinical scenario and frequency of NIC generation.

**Figure 2.**
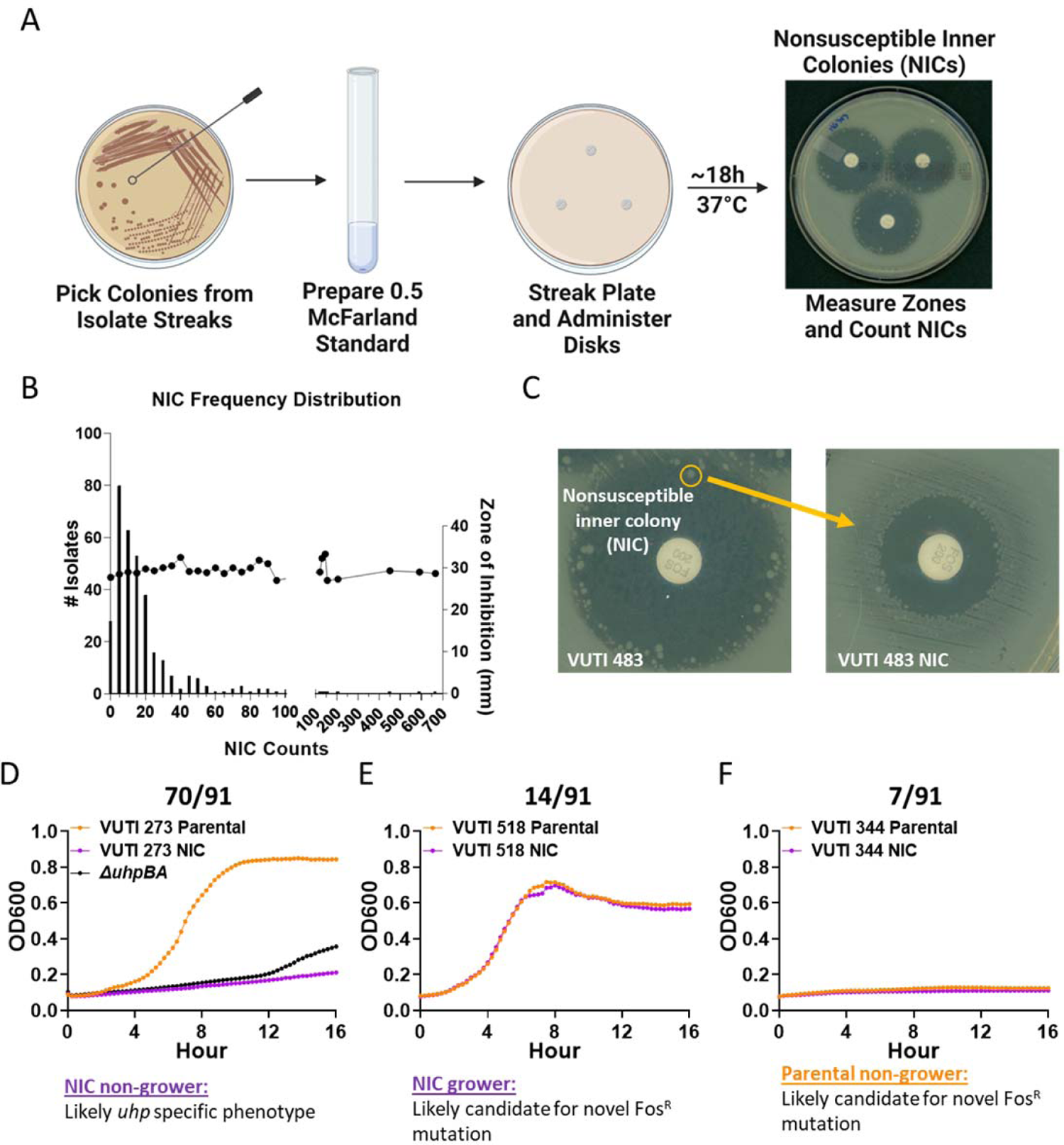
Most clinical UPEC isolates form fosfomycin non-susceptible inner colonies (NICs). **A)** Schematic depicts the Kirby-Bauer disk diffusion method of susceptibility testing used for screening of NIC phenotype. **B)** Histogram depicts the number of UPEC isolates (left y-axis) shown to generate NICs. Average NIC counts organized into bins (x-axis), and average zone of inhibition size for each bin (right y-axis) are shown. Data shown were generated from 2 biological replicates. Source data for this figure can be found in Supplementary File 1. **C)** Photo depicts the fosfomycin disk diffusion profile of a representative isolate VUTI 483 (left) and one randomly selected NIC (right). A complete profile of all the strains and NICs analyzed in this study is found in Supplementary Figure S2. **D-F)** Growth curves depict a single representative from the three different growth patterns observed for daughter NICs (purple) compared to the parent UPEC isolates (orange) during growth in M9 minimal media supplemented with 0.4% glucose-6-phosphate (G6P). A total of 70/91 NICs tested did not grow on G6P media compared to their parent strain (D), while 14/91 NICs grew in a manner similar to their parent strain (E). A third subset, 7/91 isolates (NICs and parent strains) displayed overall poor growth in G6P as a sole carbon source. Growth phenotypes were designated following duplicate biological replicates with 4 technical repeats each (Source data in Supplementary File 1).

When isolated and subsequently tested by Kirby-Bauer disc diffusion assays, the Fos^R^ NICs demonstrate significantly smaller zones of inhibition compared to the parent strain (**Fig. 2C, Fig. S2, Supplementary File 2)**. These results suggest that the fosfomycin resistance of NICs comes from stable genetic mutations. Therefore, we next sought to identify the mutations associated with fosfomycin resistance in a representative cohort of NICs.

### Novel pathways connected to fosfomycin resistance exist

Previous work showed that a common route to the NIC/Fos^R^ phenotype is through genetic disruption of the *uhp* two component system or the hexose phosphate transporter *uhpT* (**Fig. 1A, C** and [16-17]). Loss-of-function mutations in *uhpBAC* or *uhpT* abrogate the ability of *E. coli* to import G6P and thus the corresponding mutants cannot grow on media in which G6P is the sole carbon source. To assess the prevalence of *uhp* mutations in UPEC NICs, we devised a simple phenotypic assay, where NICs and their parental UPEC strains were screened with in M9 minimal media supplemented with G6P as the sole carbon source. We surmised that if the ability to transport G6P is hampered via the mutation of *uhpBA* or *uhpT*, then the corresponding NIC would not be able to import G6P and would therefore exhibit a growth defect, relative their parent strain. However, if other gene mutations confer resistance to fosfomycin, then the NIC and parent strain may exhibit similar growth on G6P as the sole carbon source. Using this scheme, we tested 91 randomly selected NIC-parent pairs, which constitute 27% of the 337 strains used in the NIC screening (**Supplementary file 2**).

Our analyses revealed 3 distinct growth patterns on G6P as the sole carbon source. Figure 2D-F depicts a representative of each of these three growth pattern, while Supplementary File 2 contains all the source data for the 91 parent-NIC pair growth curves. Out of the 91 parent-NIC pairs, 71 NICs did not grow on G6P relative to their parent, and these were deemed as “NIC non-growers” (**Fig. 2D, supplementary File 2**). Fourteen out of the 91 parent-NIC pairs displayed equally good growth on G6P (“NIC-growers, **Fig. 2E, Supplementary File 2**), while in the remaining 7 NIC-Parent pairs neither the parent, nor the NIC grew in G6P media (**Fig. 2F, Supplementary File 2**). These data suggest that loss of G6P import – likely through UhpT is indeed the most common route to fosfomycin, but it also revealed that additional genes influence fosfomycin resistance, in agreement with previous studies [16].

We then proceeded to identify genomic differences between the 91 Fos^R^ NICs and their paired parent isolates. We isolated genomic DNA from each NIC-parent pair and subjected the samples to whole-genome sequencing and subsequent variant analysis. Of 91 pairs sequenced, two were eliminated for failing quality control. The two eliminated pairs were in the “non-grower category” shown in Fig. 2F. For the remaining 89 NIC-parent pairs, identified SNPs were organized by nonsynonymous mutation based on a minimum of 30x sequencing coverage and at least 60% of reads containing the identified SNP. All parental isolate reads and assemblies are publicly available under BioProject Number PRJNA819016. All NIC-associated reads are publicly available under BioProject Number PRJNA975897.

In line with the growth profiles on G6P, 66% of NICs (59/89) harbored *uhp* mutations in *uhpB, uhpA, uhpC* or *uhpT* (**Table 1**), the majority of which are in the NIC non-grower category (**Fig. 2D**). Intriguingly, several mutations, especially in *uhpC*, were identified in NIC-growers (**Table 1**, **Fig. 2E**). These are very interesting, as UhpC constitutes the sensing component of the Uhp signaling system, bears similarity to UhpT, and has been previously shown to transport G6P albeit poorly [24]. These mutations may be allowing UhpC to transport G6P into the cell, but not fosfomycin. Of the remaining 30 NIC-parent pairs, 4 NICs harbored high-confidence, high-effect mutations in a single non-*uhp* locus (**Table 2**). The remaining 26 NIC-parent pairs harbored too many SNPs to confidently identify a single locus that contributes to fosfomycin resistance and were therefore not considered further. From the confidently identified loci (**Table 2**), we focused on investigating the effects of *pykF* that encodes an aerobic pyruvate kinase in *E. coli* (**Fig. 1D**) We selected *pykF*, because a NIC harboring a premature stop codon in *pykF* (HG320 NIC) displayed the highest NIC MIC (>1024µg/mL), which is 16-fold higher than the fosfomycin MIC of the parental isolate (**Table 2**, **Fig. S3**).

**Table 1.** Number of strains with high-confidence SNPs in *uhp* and non-uhp regions for 89 NIC-Parent pairs.

| Locus | Number of unique strains with SNPs in this locus |  |  |
| --- | --- | --- | --- |
|  | NIC non-grower | NIC grower | Parental and NIC non-growers |
| *uhpT* | 14 | 1 | 3 |
| *uhpB* | 15 | 3 | 0 |
| *uhpA* | 9 | 2 | 0 |
| *uhpC* | 7 | 5 | 1 |
| *Non-uhp* | 24 | 3 | 3 |

**Table 2.** High-confidence mutations in non-*uhp* regions, determined by high sequence coverage, and variant frequencies of at least 60% of reads.

| Strain Designation | G6P growth pattern | Fosfomycin MIC | SNP locus/Protein predicted or known function |
| --- | --- | --- | --- |
| VUTI 519 NIC | Parental and NIC non-grower | >1024µg/ml | *pykF* – Aerobic Pyruvate kinase (Fig. 1D) |
| VUTI 516 NIC | Parental and NIC non-grower | 512 µg/ml | *rdcB* – Diguanylate cyclase regulator RdcB family protein |
| VUTI 266 | NIC non-grower | 256µg/ml | *ygeK* – Response regulator that interacts with UhpB |
| VUTI 286 | NIC non-grower | 256µg/ml | *fadK* – Short chain fatty acid CoA ligase |

### Clean Deletions of known and novel genes associated with fosfomycin resistance increase the fosfomycin MIC of a susceptible *E. coli* strain

To evaluate the specific contribution of *uhp* and *pykF* in increasing the MIC of a fosfomycin susceptible strain, we created clean deletions of *uhp* genes and *pykF* in the well-characterized UPEC strain UTI89 that is fosfomycin-susceptible ([25] and **Fig. S3**). PykF and its anaerobic homolog *pykA* encode for enzymes Pyk I and Pyk II respectively, which catalyze the conversion of PEP to pyruvate and ATP (**Fig 1D**). Disruption of *pykF* may be increasing the intracellular PEP levels, outcompeting fosfomycin binding. Given the intersection of these enzymes with glycolysis and entry into the TCA cycle, we generated clean deletions of *pykF, pykA*, or both, in UTI89. The resulting markerless deletion strains exhibited substantial increases in MIC, with Δ*uhpBA* and Δ*uhpT* having an MIC 128 µg/mL compared to 1µg/mL in wild type UTI89 (**Fig. S3**). Similarly, we observed a 32-fold increase (32µg/mL) in UTI89*ΔpykF*. While loss of *pykA*, the pyruvate kinase expressed under anaerobic conditions in *E*. *coli*, did not affect the MIC of UTI89*ΔpykA* under the tested aerobic condition, deletion of both pyruvate kinases increased the MIC of UTI89*ΔpykAΔpykF* to 64µg/mL (**Fig. S3**). Together these data indicate that clean deletion of *pykF* increases the fosfomycin MIC in *E. coli* strains. We next investigated whether mutations in *uhp* or *pyk* impart a fitness cost in during UTI.

### Loss of *uhp* genes – the most common route to Fos^R^ – is not associated with fitness costs during urinary tract infection

Previous studies correlated the increased Fos^R^ in NICs to mutations within the *glpT* transporter gene, as well as components of the UhpBA two-component system and its regulon, which affect fosfomycin import into the cell [11], [14], [16]. Indeed, deletion of the two-component system *uhpBA* that positively regulates *uhpT* in cystitis isolate UTI89 renders the resulting strain fosfomycin resistant (**Fig. S3**). In limited studies, NICs with mutations in *uhp* genes were shown to exhibit decreased growth rates *in vitro*, in laboratory media and in urine culture [16-17]. These previous reports contributed to the current assumption that mutations associated with fosfomycin resistance are broadly accompanied by a high biological cost. This assumption has led to the differences in fosfomycin susceptibility testing protocols between the US (CLSI) and European (EUCAST) clinical laboratory settings, where EUCAST guidelines call for ignoring NICs, while CLSI calls for taking them into account [15]. This creates confusion in the clinical laboratory setting globally, further complicating efforts for antibiotic stewardship. Previous studies also reported that mutations in the *uhp* system impair UPEC colonization of the host kidney in a UTI model [17], however no data were provided regarding bladder colonization, chronic infection, or long-term persistence in reservoir niches, which is a hallmark for recurrent infection with UPEC [26-27].

To gauge the contribution of the UhpBA two-component system during pathogenesis, we leveraged our expertise in well-established murine models of UTI [28]–[30] to evaluate the fitness of *ΔuhpBA* compared to its isogenic parent during acute and chronic UTI. Moreover, we measured the ability of each strain to transit to and persist in the gut following a UTI.

Prior to introducing the strains into animals, we wanted to determine if a growth defect exists during growth in urine. Strains were therefore grown in pooled urine under hypoxic conditions to mimic the host urinary tract and evaluate the presence of any apparent growth defect. Following 8h of growth, no significant difference in growth is detected between *ΔuhpBA* and wild type UTI89 (**Fig 3A**). Strains were then grown and prepared for inoculating 7-8 week old female C3H/HeN mice as we previously described [29]. Following transurethral inoculation, infected mice were euthanized 16 hours post-infection (hpi) to assess whole bladders and kidneys for CFUs at the acute infection stage during the time when intracellular biofilm-like communities (IBCs) form. These experiments revealed no significant differences in bladder or kidney titers between Δ*uhpBA*- and wild type-infected mice (**Fig. 3B-C**). To visualize the ability of Δ*uhpBA* to form IBCs, we used strains that were transformed with plasmid pANT4 that constitutively expresses GFP. We detected no significant differences in the number (**Fig. S4A**) or morphology (**Fig. S4B**) of IBCs formed by the wild-type (WT) and Δ*uhpBA.* These data indicate that deletion of *uhpBA* does not impart a fitness defect during acute UTI.

**Fig. 3.**
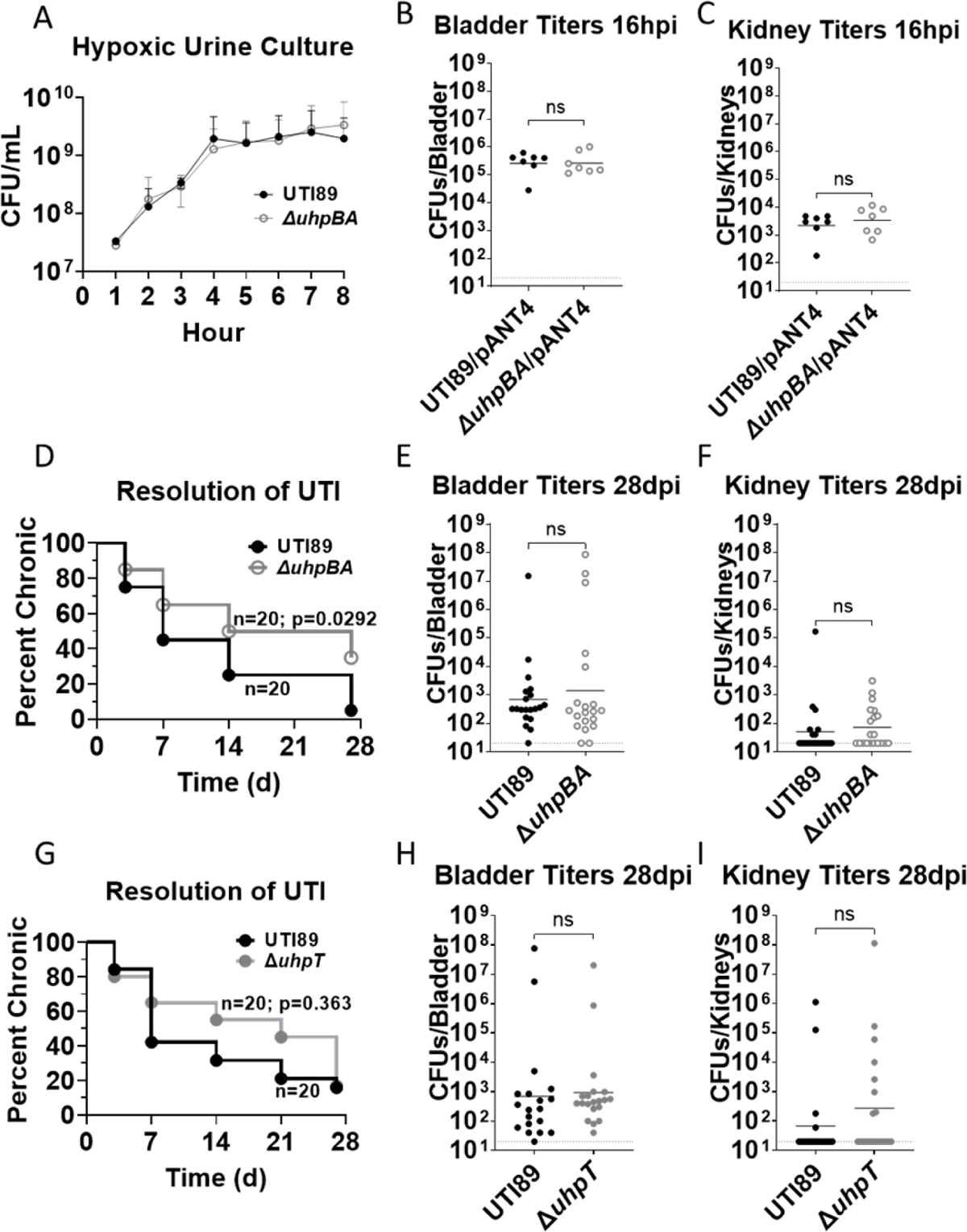
U*h*p mutants do not display fitness defects during acute or chronic UTI. **A)** Graph depicts growth of UTI89 (black) and Δ*uhpBA* (grey) in pooled urine under 4% O_2_ that mimics the host environment. The average of 3 biological repeats is shown. **B-C)** Graphs depict organ titers at 16 hours post infection (hpi) for bladders **(B)** or kidneys **(C)** from mice infected with UTI89 (black) or Δ*uhpBA* (grey). Statistical comparisons were performed in Graphpad using a two-tailed Mann-Whitney. **D)** Kaplan-Meier plot depicts the rate of bacteriuria resolution in mice infected with Δ*uhpBA* (grey) compared to mice infected with UTI89 (black), over a period of 28 days. **E-F)** Bladder **(E)** and kidney **(F)** titers at 28 days post infection (dpi) in mice infected with UTI89 (black) or Δ*uhpBA* (grey) populations. Statistical comparisons were performed in Graphpad using a two-tailed Mann-Whitney. **G)** Kaplan-Meier plot following resolution of bacteriuria in mice infected with Δ*uhpT* (open circles) compared to UTI89 (black) over 28 days. **H-I)** Bladder **(H)** and kidney **(I)** titers at 28 dpi in mice infected with UTI89 (black) or Δ*uhpT* (open circles) populations. Statistical comparisons were performed in Graphpad using a two-tailed Mann-Whitney.

We next repeated the transurethral inoculations, but this time we monitored infection progression over a 28-day period. In C3H/HeN mice, chronic infection is characterized by persistent bacteriuria, where greater than 10,000CFU/ml of urine are shed daily be infected animals [26]. Using longitudinal urinalysis, we observed that Δ*uhpBA*-infected mice resolve bacteriuria more slowly than the isogenic parent (**Fig. 3D**) and organ titers at 28dpi are indistinguishable from those of the parent strain (**Fig. 3E, F**). Consistent with slow resolution, we see high numbers of bacteria in the gut, following infection (**Fig. S5**). These results demonstrate that Δ*uhpBA* does not exhibit a fitness defect *in vivo* and can colonize infection- and reservoir niches as efficiently as the WT parent.

Given that UhpBA signaling leads to the upregulation of the UhpT transporter, we also evaluated the fitness of a mutant deleted for the *uhpT* gene. Similar to the observations with Δ*uhpBA*, Δ*uhpT* exhibits no acute or chronic colonization defect (**Fig. 3G-I**). However, the Δ*uhpT* mutant has similar resolution rates as the isogenic parent (**Fig. 3G**) suggesting that in UPEC, UhpBA may have a more expanded regulon that extends beyond *uhpT* and may be responsible for the increased persistence of bacteriuria observed in Δ*uhpBA*. Combined, these data indicate that mutation of the most common route to fosfomycin resistance is not associated with a fitness defect in the bladder.

### Loss of *pykF* – a newly identified route to Fos^R^ in *E. coli*– does not come at a fitness cost to UPEC

Previous studies of UPEC pathogenesis showed that steps in glycolysis to be expendable during acute infection [18-19] Our work demonstrating that UPEC respires aerobically during infection [31-32] combined with further work indicating amino acid utilization during infection [18-19] we hypothesized that single loss of *pykA* (Pyk II) or *pykF* (Pyk I) would not come at a fitness cost during infection. We thus took a similar approach to test the colonization potential of Δ*pykF*, Δ*pykA* and Δ*pykA*Δ*pykF* during mono-infections using the murine models discussed above.

Given the role of the Pyk enzymes in energy metabolism and our prior observations indicating UPEC aerobically respires inside the urothelial cell [29], we wanted to evaluate the role of each *pyk* in UPEC intracellular expansion. We thus incorporated an additional time-point, 6h post infection, which marks the early development of IBCs and allows us to determine how many bacteria are intracellular by treating the excised bladders with gentamicin [29]. These experiments revealed that during acute infection, the numbers of whole bladder organ (**Fig. 4A**) or intracellular (**Fig. 4B**) CFUs are comparable in wild-type UTI89, Δ*pykA*, Δ*pykF*, and Δ*pykA*Δ*pykF*. Similarly, all strains had comparable kidney titers at 6h post infection (**Fig. 4C**). Strains continued to have comparable organ titers at 24hpi (**Fig. 4D, E**).

**Fig. 4.**
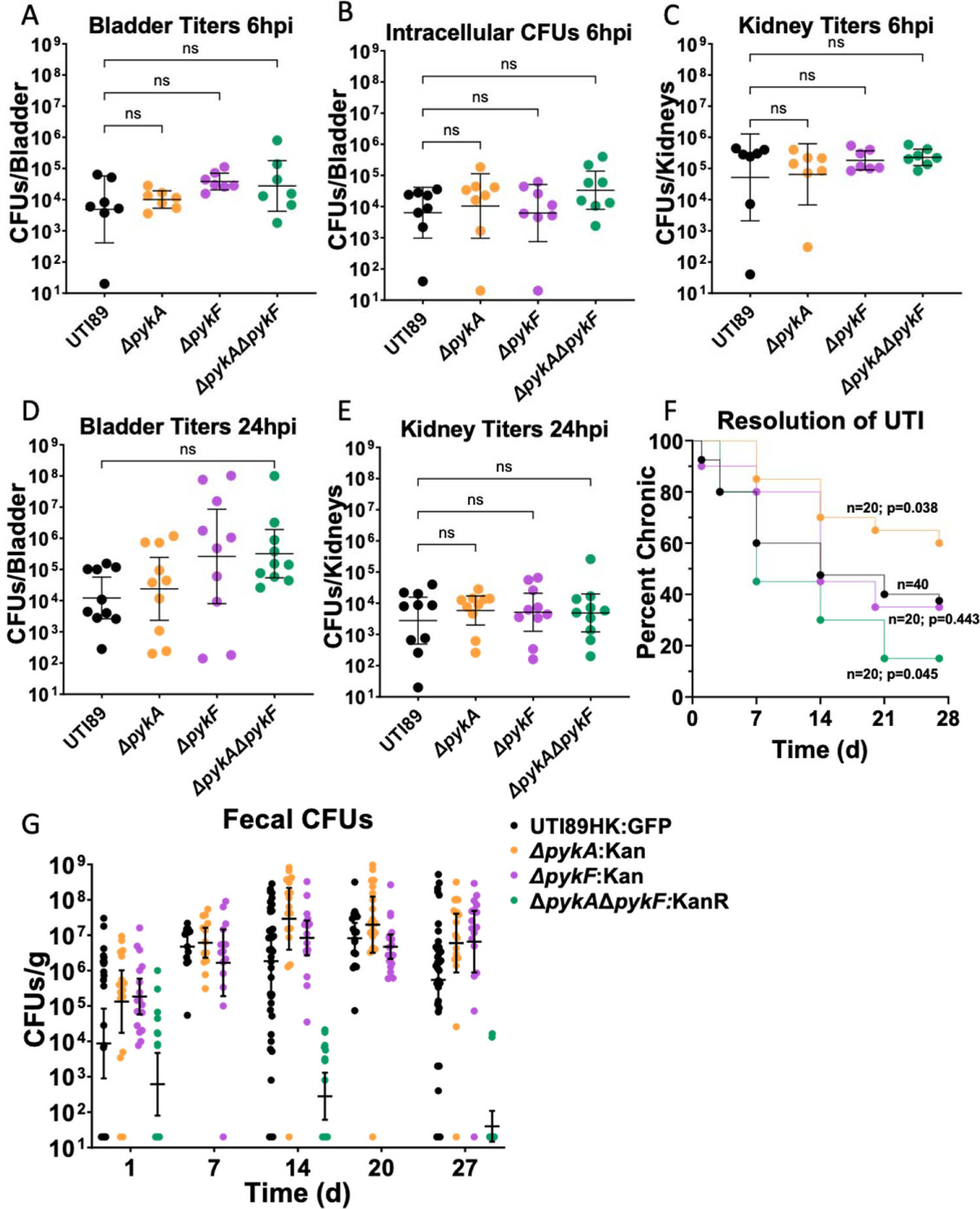
Evaluation of *pyk* mutant fitness *in vivo*. **A-B**) Graphs depict total (A) or intracellular (B) bladder titers for wild-type UTI89 (black), Δ*pykA* (orange), Δ*pykF* (purple) or the double mutant ΔpykAΔ*pykF* (green) at 6hpi. **C)** Graph depicts kidney titers for each strain at 6hpi. **D-E)** Graphs depict total bladder (D) or kidney (E) titers 24hpi. Statistical comparisons of each strain with the wild type parent for panels A-E were performed using two-tailed Mann Whitney within Graphpad software. **F)** Kaplan-Meier plot depicts the time to resolution of bacteriuria over time in mice infected with wild-type UTI89 (black), Δ*pykA* (orange), Δ*pykF* (purple) or the double mutant ΔpykAΔ*pykF* (green) over a period of 28 days. Statistical analysis was performed in Graphpad Prism using the Mantel-Cox test. **G)** Graph depicts fecal CFUs over time. Fecal samples were collected contemporaneously with the urinalysis in panel F.

We next monitored infection progression over a 28-day period. Using longitudinal urinalysis, we observed that Δ*pykF*-infected mice resolved bacteriuria equivalently to mice infected by UTI89 (**Fig. 4F**). However, the mutant deleted for the anaerobic pyruvate kinase *pykA* resolved bacteriuria more slowly (**Fig. 4F**). This was accompanied with Δ*pykF*-infected mice demonstrating significantly higher total bladder CFUs on the 28th-day post infection (**Fig. S6A**) This observation is in agreement with previous studies by Alteri *et al.*, who showed a competitive advantage of Δ*pykA* during co-infection studies in a different UPEC strain [19]. Conversely, while the double mutant Δ*pykA*Δ*pykF* displays no virulence defects during acute infection, mice infected with this mutant resolved bacteriuria more rapidly (**Fig. 4F**), had significantly lower fecal titers over time (**Fig. 4G**), and demonstrated significantly lower total bladder and kidney CFUs at 28 dpi (**Fig. S6C, D**). Combined, these data indicate that at least one new route to fosfomycin resistance in UPEC – inactivation of PykF– does not impair virulence and uncover that PykA and PykF are functionally redundant for pathogen persistence.

## Discussion

In this work, we demonstrate that several routes to fosfomycin resistance exist and that at least two of these routes come at no fitness cost to the uropathogen. We show, utilizing a representative UPEC strain cohort, that most clinical isolates produce non-susceptible inner colonies (NICs) during fosfomycin AST. This phenomenon is well established, but largely attributed as an artifact of *in vitro* testing. Moreover, NICs have previously been associated with high biological cost in limited characterization studies [15-17]. In this paper, we present the first study, to our knowledge, to critically evaluate colonization and persistence of NIC-associated mutations within the host. We believe that thorough understanding of the mechanisms leading to fosfomycin resistance is of utmost importance as the use of this antibiotic continues to increase.

Previous work pointed to *uhp* system mutations as the most common route to the NIC phenotype [14], [16] and concluded that the inability to transport a key glycolysis intermediate, G6P, would impose substantial metabolic stress to the pathogen. We and others have demonstrated that glycolysis is dispensable in the bladder environment, where glucose and intermediates thereof is scarce. This work demonstrates that UPEC can persist in the gut without the need to import G6P, or without one of the two pyruvate kinases it encodes, but not both. Therefore, the notion that emergence of fosfomycin resistance is compounded by the need to perform glycolysis is not necessarily true.

We used growth on G6P as the sole carbon source as a quick method for screening in search of NICs that may not have mutations in *uhp*. NICs that failed to grow in M9 media supplemented with G6P (NIC non-growers), were mostly associated with *uhpBA* mutations, while NICs that successfully grew in G6P harbored – as we showed – non-*uhp* routes to Fos^R^. Surprisingly, a third growth phenotype was observed in our G6P growth assays: in which neither the NIC nor its parent strain grew in G6P media. This phenotype suggested either absence of *uhp* genes from the parent strain, or the presence of loss-of-function mutations in the parental *uhp* genes. The complete absence of *uhp* genes was confirmed in clinical isolate VUTI 516, while the remaining parent non-grower strains harbored multiple SNPs in their corresponding *uhp* loci. This is an important observation, since tested UPEC strains were acquired directly from patients at Vanderbilt University Medical Center, who were not prescribed fosfomycin for treatment, yet they harbored fosfomycin resistant isolates that may have arisen in response to an unknown stress *in vivo*. These representative strains demonstrate that despite the absence of a functional *uhp*, they established infection in humans. Additionally, this further supports the need for a more critical evaluation of strains containing NIC-associated mutations, which have previously been deemed of high biological cost to the pathogen. Another critical mutation identified in a parental and NIC non-grower was within the *pykF* gene, again indicating that in the host, UPEC does not rely on glycolysis [18].

Among identified NIC mutations, *uhp* system mutations were the most abundant associated route with NIC Fos^R^. It should be noted however, that among 5/14 NICs that grew in the presence of G6P harbored mutations within the *uhpC* sensor gene. Such mutations may be altering the function of UhpC (that bears sequence similarity to UhpT) such that it does not induce the activation of UhpBA signaling (**Fig. 1A**) but allows UhpC to serve as a G6P transporter. We are currently investigating how mutations in *uhpC* alter its signaling capacity.

In limited studies, NICs with mutations in *uhp* genes had been shown to exhibit decreased growth rates *in vitro*, in laboratory media and in urine culture [16-17]. These initial characterizations contributed to the belief that mutationally acquired fosfomycin resistance is broadly accompanied by a high biological cost. Currently, this has contributed to differences in US (CLSI) and European (EUCAST) clinical standards during fosfomycin testing in which EUCAST guidelines call for ignoring NICs, while CLSI calls for taking them into account [15]. However, through a thorough evaluation of acute and chronic stages of infection, we have demonstrated that both common and uncommon routes to fosfomycin nonsusceptibility fail to negatively impact colonization and persistence within the host urinary tract. Based on our data, we raise the alarm that multiple routes that lead to fosfomycin resistance do not affect pathogenesis and propose that EUCAST and CLSI guidelines reevaluate how they classify fosfomycin resistant subpopulations during susceptibility testing.

## METHODS

### Strains, plasmids and primers used in this study

Strains used in this study are listed in Supplementary file 1a, along with associated de-identified patient information. The well-characterized cystitis isolate UTI89 was used as a reference strain in which clean deletions were created. Strains and primers used in these studies are found in **Supplementary File 1.**

### Kirby-Bauer Disk Diffusions

Susceptibility testing of 337 UPEC clinical isolates from VUMC’s microVU biorepository (https://www.vumc.org/microvu/home) was performed in accordance with established Kirby Bauer disk diffusion protocols [33]. (MicroVU banks an average of 13,000 UPEC isolates. Power analysis was performed to determine the number of isolates required to estimate the average NIC count of 13,000 annual *micro*VU UPEC isolates with a margin of error of 5 and a confidence interval of 95%). Each UPEC isolate was streaked for isolation. Single colonies were picked and suspended in sterile saline to match to a 0.5 McFarland Standard. Suspensions were used to inoculate nonselective Mueller-Hinton agar plates. Three fosfomycin disks (BD BBL) were placed equidistant on each plate. Nonsusceptible inner colonies (NICs) were counted, and zones of inhibition were recorded in accordance with EUCAST guidelines. Experiments were repeated at least twice for each isolate. Representative NICs were sub-cultured and freezer stocks were prepared for subsequent sequencing analysis. The same protocol was followed for ETEST® strips (bioMérieux), but with one E test strip added to the center of the Mueller-Hinton plate, instead of three discs.

### Growth conditions

All strains were propagated in lysogeny broth (LB) at 37°C with shaking for routine culture and cloning or gene deletion purposes. <u>For the G6P growth analyses</u>: Growth curves of parental UPEC isolates and their respective NICs were performed in M9 media [30] supplemented with 0.4% G6P as the sole carbon source. The *ΔuhpBA* mutant strain served as a negative growth control. Clinical isolates and lab strains were streaked for isolation on LB agar and incubated at 37°C overnight. Cultures were created in 5mL LB from single colonies of each parental and paired NIC isolate and incubated at 37°C shaking overnight. Strains were sub-cultured in 5 mL of fresh LB in a 1:1000 ratio for 2 hours, or until growth was observed. Each subculture was inoculated into M9 media supplemented with 0.4% G6P in a 1:100 ratio. Strains were loaded in 96 well plates and absorbance was measured at 600nm for 15 minute intervals over 16 hours of growth with continuous orbital shaking. Instances in which NIC growth was observed were subsequently validated with further repeats. <u>For animal studies:</u> Bacteria were first propagated in LB with shaking overnight. Overnight cultures were used to seed fresh LB at a ratio of 1:1000 and incubated statically for 24h. Static cultures were passaged one more time in LB at a 1:1000 ratio and incubated statically for another 24h prior to preparing for inoculation in mice. This 2 x 24h static culture has been previously shown to enhance expression of type 1 pili which are adhesive fibers required for the initiation of infection [34].

### Next Generation Sequencing and Analysis

Clinical UPEC isolates and their paired representative NICs were streaked for isolation and propagated in overnight cultures as described above. Aliquots of 750µL of overnight were pelleted by centrifugation and treated for genomic DNA isolation with the PureLink™ Genomic DNA Kit. Parental isolates were prepared for sequencing using the Illumina DNA Prep Kit (catalog no. 20018705). These isolates were sequenced on an Illumina MiSeq or NextSeq 2000 instrument producing 2×151 reads. The parental isolate reads were evaluated for quality using FastQC v. 0.11.9 with default options. We assessed the read files for quality by looking at the number of sequences with over 85% Q30 base pair reads and overall passing all checks for base quality, GC content, N content, length distribution, and overrepresented sequences [35]. Reads were trimmed using TrimGalore v. 0.6.7, with option –paired. The reads were assembled *de novo* using SPAdes v. 1.13 with options –isolate and for paired reads. The resulting FASTA file was finally filtered and contigs <800bp were removed. Code for trimming and assembly of the parental isolates can be found at www.github.com/gracehm. Parental FASTA files were annotated using the workspace on Bacterial and Viral Bioinformatics Resource Center (BV-BRC) v. 3.30.5i using the RAST tool kit [34-35]. NIC read files were mapped against their parental FASTA files using BWA-mem through BV-BRC [38]. Variants in the NICs were identified using FreeBayes [39]. Variants were assessed for potential SNPs eliciting their increased Fos^R^ phenotype. SNPs found within coding sequences were prioritized based on a frequency of reads >60%.

### Data Availability

All sequencing data produced from this work were deposited in the NCBI Sequencing Read Archive (SRA) and Genbank. All parental isolate reads and assemblies are associated with BioProject Number PRJNA819016. All NIC associated reads are associated with BioProject Number PRJNA975897.

### Murine Infections

Murine infections were performed as previously described [34]. In brief, UTI89 and isogenic mutants were grown overnight in 5 mL LB medium and grown while shaking at 37°C. Cultures were then diluted 1: 1000 into 10 mL fresh medium and grown at 37°C statically for 24h. Cultures were subsequently diluted 1: 1000 into 10 mL fresh medium and grown again at 37°C statically for 24h. Cultures were normalized in sterile PBS and 7- to 8-week-old C3H/HeN female mice were transurethrally inoculated with 50 µL containing 10^7^ CFUs of bacteria. Mice were sacrificed at 6, 24h or 28 days post infection, in which bladders and kidneys were removed, homogenized with an Omni tissue homogenizer, and serial diluted for CFU enumeration. Intracellular CFUs were quantified following PBS wash of the bladders and subsequent treatment with 100µg/mL gentamicin in PBS for 90 minutes. After gentamicin treatment, bladders were washed in PBS, and homogenized in PBS containing 0.1% Triton X-100 before serial dilution and plating. Animal studies were approved by the Vanderbilt University Medical Center Institutional Animal Care and Use Committee (IACUC) (protocol number # M1800101-01).

